# Dissection of the microProtein miP1 floral repressor complex in Arabidopsis

**DOI:** 10.1101/258228

**Authors:** Vandasue L. Rodrigues, Ulla Dolde, Daniel Straub, Tenai Eguen, Esther Botterweg-Paredes, Bin Sun, Shinyoung Hong, Moritz Graeff, Man-Wah Li, Joshua M. Gendron, Stephan Wenkel

## Abstract

MicroProteins have emerged as potent regulators of transcription factor activity. Here we use a combination of forward genetics and proteomics to dissect the miP1a/b microProtein complex that acts to delay the floral transition in Arabidopsis. The microProteins miP1a and miP1b can bridge an interaction between the flowering promoting factor CONSTANS (CO) and the TOPLESS (TPL) co-repressor protein to represses flowering. We find that the JUMONJI14 (JMJ14) histone demethylase is part of this repressor complex that can initiate chromatin changes in *FLOWERING LOCUS T* (*FT*) gene, the direct target of CO. Plants with mutations in *JMJ14* exhibit an early flowering phenotype that is largely dependent on the activity of *CO*, supporting a role for CO in this repressive complex. When mis-expressed at the shoot apex, *CO* can induce early flowering only in the *jmj14* background. Our results indicate that the repressor acts in the shoot apical meristem to keep it in an undifferentiated state until the leaf-derived florigen signal induces the conversion into a floral meristem.

## INTRODUCTION

Annual plants, such as the model plant *Arabidopsis thaliana*, induce flowering only once during their lifetime and this transition occurs in a non-reversible fashion. Once a plant commits to flowering it cannot return back to the vegetative growth stage. In order to maximize reproductive success, the plant integrates seasonal information such as temperature and day-length to initiate flowering only under the most optimal conditions. Arabidopsis being a long-day plant will start flowering when days are long, thus restricting flowering to occur in summer.

The molecular network underpinning the photoperiodic flowering response has been elucidated using mutants and ecotypes showing variations in their respective flowering phenotypes (Andres and Coupland, 2012; Imaizumi and Kay, 2006). A central component of the photoperiodic flowering time pathway is the CONSTANS (CO) transcription factor (Putterill et al., 1995). Both *CO* mRNA and protein exhibit diurnal patterns of expression but the protein can only accumulate at the end of long days (Suarez-Lopez et al., 2001; Valverde et al., 2004). Once CO protein is present, it acts as a transcriptional activator and induces expression of *FLOWERING LOCUS T* (*FT*) (Samach et al., 2000). Both CO stability and its interaction with the *FT* promoter are mediated by a set of PSEUDO RESPONSE REGULATOR (PRR) proteins (Hayama et al., 2017).

Arabidopsis leaves act as photoperiod sensors and both *CO* and *FT* are expressed and active in the leaf vasculature. When expressed from a phloem-specific promoter, CO is able to fully rescue the late flowering phenotype of *co* loss-of-function mutant plants while expression in the shoot apical meristem does not complement the late flowering (An et al., 2004). This contrasts the finding that expression of *FT* in either the shoot meristem or the leaf vasculature is effective in triggering an early flowering response (An et al., 2004). Later, it was revealed that CO acts in the phloem to induce *FT* expression and the resulting FT protein acts as a systemic florigen signal that travels from the leaves to the shoot meristem where it initiates the conversion of the vegetative leaf-producing meristem into a reproductive flower-producing meristem (Corbesier et al., 2007; Jaeger and Wigge, 2007; Mathieu et al., 2007; Tamaki et al., 2007).

Recently, we identified two Arabidopsis microProteins, miP1a and miP1b, that can interact with CO and when overexpressed cause a late flowering phenotype (Graeff et al., 2016). MicroProteins are small single-domain proteins that can exist as individual genes in the genomes of higher eukaryotes. MicroProteins are sequence-related to larger, multi-domain proteins and have evolved during genome evolution by amplification and subsequent degeneration. A hallmark of microProtein function is the presence of a single protein domain, often a protein-protein-interaction domain, allowing the microProtein to exert dominant-negative modes of action by sequestering target proteins (Eguen et al., 2015; Staudt and Wenkel, 2011). In the case of miP1a/b it is however not a simple sequestration but rather the formation of a higher order repressor complex (Graeff et al., 2016). In this repressor complex, the microProteins bridge CO and the TOPLESS (TPL) co-repressor protein. Given the significant role of TPL in this repressor complex, we reasoned that other accessory proteins might be involved. In order to identify such partners, we carried out EMS mutagenesis with plants ectopically expressing *miP1a* that flower very late. In this EMS screen, we identified *SUPPRESSOR OF MIP1A-1* (*SUM1*), having a frame-shift mutation in the Histone 3 Lysine 4 (H3K4) -demethylase *JUMONJI14* (*JMJ14*), causing *miP1a*-overexpressing plants to flower early.

*JMJ14* encoding a H3K4-demethylase has a known role in the regulation of flowering (Yang et al., 2010). H3K4 methylation is associated with active chromatin, hence removing H3K4 methylation acts to initiate gene silencing. Plants carrying loss-of-function mutations in *JMJ14* display an early flowering phenotype, express higher levels of *FT* and have increased levels of H3K4 methylation in the *FT* promoter (Yang et al., 2010). JMJ14 plays an additional role in RNA silencing and has been shown to also influence DNA methylation in the process of silencing transposon transcripts (Searle et al., 2010).

In this study, we identify JMJ14 as a component of the microProtein floral repressor complex. Mutations in *JMJ14* suppress the late flowering phenotype exerted by ectopic expression of both miP1a and miP1b but cannot complement late flowering *co* mutants, indicating that JMJ14 does not act independently of CO. Additional proteomic studies revealed that JMJ14 can directly interact with TPL/TOPLESS-related (TPR) proteins. Thus, it seems that miP1a/b assemble in a larger chromatin silencing complex which likely induces chromatin changes in *FT* promoter when miP1a/b are expressed at high levels. The finding that CO is expressed in the shoot apical meristem (SAM) but cannot induce flowering when expressed there, suggested to us that CO might attain a SAM-specific function that depends on interacting partners. Both miP1a/b and JMJ14 are co-expressed with CO in the SAM where they could form a repressor complex. Exploring this hypothesis, we mis-expressed CO in the SAM of a *jmj14* mutant and observed a strong early flowering phenotype. Taken together, our findings indicate that the *FT* gene is actively repressed in the shoot apex by a repressor complex likely involving CONSTANS/ CONSTANS-like transcription factors, microProteins miP1a/b, TPL and JMJ14. This repressor complex prevents flowering until the leaf-derived FT protein triggers the transition to the reproductive growth phase.

## RESULTS AND DISCUSSION

### The microProtein repressor complex requires JMJ14 activity

MiP1a/b-type microProteins interact with CO through their B-Box domain and with TPL via a five amino-acid stretch at the carboxy-terminal end (Graeff et al., 2016). The finding that TPL is required for the microProteins to exert their strong repressive potential points towards the existence of a higher order repressor complex. In order to identify novel components of such a repressor, we performed an EMS suppressor mutagenesis with plants ectopically expressing FLAG-miP1a. In total, we identified 25 potential suppressor mutants of which four mutants no longer expressed the transgene, 13 showed expression levels between wild type and the *FLAG-miP1a* overexpressor and eight plants showed expression levels comparable to the parental plants (Suppl. Fig. S1). One suppressor, named *sum1*, was isolated and studied in detail. Under long day conditions, Col-0 wild type plants flower rapidly producing only a small number of rosette leaves, in contrast to transgenic plants over-expressing FLAG-miP1a (Fig. 1A,B) which are late flowering and produce many leaves before transitioning to flowering. Plants that are homozygote for the *sum1* mutation flower slightly earlier than wild type plants, despite the presence of the FLAG-miP1a transgene. High level of transgene expression does not always correlate with high translation. In order to determine the protein expression of miP1a, we measured the levels of FLAG-miP1a protein in the parental transgenic plant and in the *sum1* background. We detect slightly lower levels of FLAG-miP1a in the *sum1* mutant compared to wild type, but the protein is still highly abundant (Fig. 1C). These findings indicate that the factor encoded by *SUM1* is required for the miP1a microProtein to repress flowering.

**Fig. 1.**
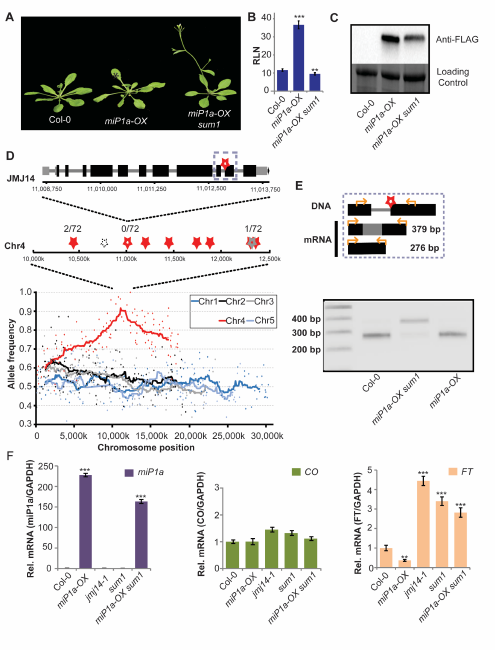
Characterization of the sum1 mutant phenotype. **(A)** Phenotype of the M2 suppressor mutant *sum1* in the *miP1a-OX* background compared to the Col-0 wildtype and the **miP1a-OX** progenitor respectively. **(B)** Determination of flowering time by counting the number of rosette leaves at bolting. Significant differences in leaf numbers between the *miP1a-OX sum1* line compared to *miP1a-OX* progenitor were observed. P<0.01. **(C)** Analysis of protein levels of FLAG-tagged miP1a protein in the *miP1a-OX sum1* line compared to *miP1a-OX*. No signal was detected in wildtype. Coomassie staining is shown as loading control. **(D)** Top: *JMJ14* gene model with G to A single nucleotide polymorphism (SNP) in the splice site before the second exon (black box: exon, grey: UTR, star: SNP). Middle: Stars indicate SNPs in a mapping interval of chromosome 4 (filled red: CDS, open red: splice site, grey: UTR, dashed: intergenic) with observed wildtype allele frequency numbers in F2 backcross generation (total 36 plants) by Restriction fragment length polymorphism (RFLP) and/or Amplification-refractory mutation system (ARMS). Bottom: Five Arabidopsis chromosomes with allele frequency of background corrected SNPs measured by mapping-by-sequencing. *(E)* Top: model of the intron retained product (379 bp) and the correctly spliced product (276 bp mRNA). Bottom: PCR amplification of the cDNA from Col-0 wild type, *miP1a-OX sum1* and *miP1a-OX* lines. *MiP1a-OX sum1* shows a predominantly higher product indicating intron retention due to a splicing defect. **(F)** Quantification by qRT-PCR of *miP1a*, *CO* and *FT* levels. *MiP1a* transgene levels are expressed at a high level in *miP1a-OX sum1* line. While *CO* levels are not significantly different, there is a significant increase in the *FT* levels in the *miP1a-OX sum1* compared to *miP1a-OX*. **P<0.005, ***P<0.001.

We next aimed to identify the causal mutation underlying the sum1 phenotype. Therefore, we isolated DNA from 20 *sum1* suppressor mutants out of a segregating F2 population from a back-cross to Col-0. All 20 individuals showed resistance to the herbicide BASTA, had high levels of *miP1a* mRNA and showed an early flowering phenotype. Whole genome sequencing of this pool of suppressor mutants and the respective parental plant identified 591 EMS-induced SNPs with a strong frequency enrichment in the middle of chromosome 4 (Supplementary dataset 1). At the summit region of the peak, we identified a mutation in the *JMJ14* gene affecting a splice junction (Fig. 1D). Further characterization of additional 36 segregating suppressor mutants revealed that all 72 examined chromosomes carried the *jmj14* mutation while flanking mutations were still segregating. We then tested if the identified mutation would interfere with correct splicing of the *JMJ14* transcript. RT-PCR-amplifications spanning the intron in question using cDNAs prepared from Col-0, transgenic *miP1a-OX* and *miP1a-OX sum1* plants revealed intron-retention in the *sum1* background (Fig. 1E). The retained intron results in a premature stop-codon and the resultant mutated JMJ14 protein lacks the carboxy-terminal FY-rich (FRYC) domain that might engage in protein-protein-interactions (Pless et al., 2011). To confirm that *sum1* is indeed the causal mutation that suppresses miP1a function, we crossed homozygote *miP1a-OX sum1* plants with either homozygote *jmj14-1* or *jmj14-3* mutant plants. Both cases showed the resultant nullizygote offspring had an early flowering phenotype (Suppl. Fig. S2), which confirms that *JMJ14* is the causal gene and encodes the protein likely required for the floral repression imposed by ectopic miP1a expression.

Additional gene expression profiling experiments revealed that *miP1a* mRNA levels are highly upregulated in transgenic *miP1a-OX* and *miP1a-OX sum1* plants. CO mRNA levels are slightly upregulated in *jmj14-1* and *sum-1* mutant plants while *FT* mRNA is highly abundant in *jmj14-1*, *sum-1* and *miP1a-OX sum1* plants, explaining the early flowering behavior (Fig. 1F).

In summary, our results show that *FT* is under a constant repression by a JMJ14-containing silencing complex. Plants lacking JMJ14, show a slightly early flowering behavior in long day conditions that can be attributed to de-repression of *FT*. The observation that ectopic microProtein expression is unable to repress *FT* in a *jmj14* mutant background suggests that miP1a is either part of the repressor complex or acts upstream of the JMJ14-induced floral repression pathway. A recent study suggests that mutations in *JMJ14* can result in a re-activation of genomic regions that have undergone post-transcriptional gene silencing and additionally can decrease the expression of transgenes by affecting the chromatin of the transgene (Le Masson et al., 2012). However, we think that this is not the case in our study because *miP1a-OX sum1 (jmj14)* plants exhibit earlier flowering than wildtype and not an intermediate flowering response. In addition, these *miP1a-OX sum1* transgenic plants are fully resistant to the herbicide BASTA and the double-heterozygote from the back-cross to wildtype revealed a very late flowering phenotype.

### JMJ14 controls flowering in a CO-dependent manner

The loss of JMJ14 function disables miP1a ability to repress flowering. MiP1b is closely related to miP1a and can also strongly repress flowering when expressed at high levels (Graeff et al., 2016). To test if the observed suppressor phenotype is specific to miP1a, we crossed late flowering *miP1b-OX* plants into the *jmj14-1* mutant. Transgenic *miP1b-OX* plants homozygote for *jmj14-1* also showed an early flowering phenotype indicating that both miP1a and miP1b need JMJ14 to execute their repressive potential (Fig. 2A,B). However, when we crossed *jmj14-1* into a *co* null mutant (*co-SAIL*), we did not observe a strong complementation of the late flowering phenotype of *co* and *co jmj14-1* double mutants flowered only slightly earlier compared to *co* (Fig. 2C,D). This demonstrates that JMJ14 is required for the post-translational inhibition of CO function or its integration into a repressor complex. Loss of JMJ14 function attenuates the repressor complex, hence plants flower early. The finding that mutations in *jmj14* do not fully complement the late flowering phenotype of *co* mutants implies that JMJ14 does not operate independently of CO. In agreement with this, we find no strong promotion of flowering in *jmj14* mutants when grown under short days, a condition where CO is not active (Supplementary Fig. S3).

**Fig. 2.**
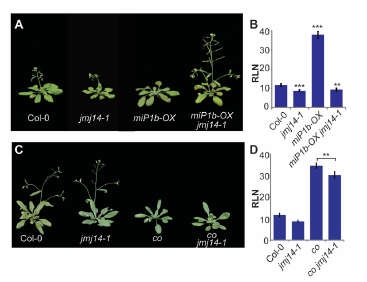
Genetic analysis of crosses between *jmj14* and other late flowering mutants. **(A)** Flowering phenotype of *jmj14-1* mutants crossed into transgenic *miP1b-OX* plants. Double mutants flower early. **(B)** Determination of flowering time by counting the number of rosette leaves at the bolting stage of wild type, jmj14-1, miP1b-OX and *miP1b-OX jmj14-1* plants. ***P<0.001. **(C)** Phenotype of *jmj14-1* introduced into *co* background is extremely late flowering compared to the parent *jmj14-1* and Col-0 wildtype. **(D)** Determination of flowering time by counting the number of rosette leaves at the bolting stage of wild type, *jmj14-1*, *co* and *co jmj14-1* mutant plants. **P<0.005, ***P<0.001.

### Dissection of the microProtein repressor complex by mass spectrometry

We previously showed that miP1a/b microProteins physically interact with CO and the corepressor protein TOPLESS (Graeff et al., 2016). Not only do miP1a/b interact with CO and TPL proteins but they act as a bridge, indicating they form an at least trimeric repressor complex. To identify additional members of this microProtein-repressor complex, we performed affinity-purification mass-spectrometry with transgenic plants overexpressing FLAG-miP1a and FLAG-miP1b (Fig. 3A,B). To identify false-positive interactors, we performed additional immunoprecipitations with non-transgenic wild type plants and plants overexpressing FLAG-GFP protein. After subtracting non-specific interactors, we identified 886 proteins interacting with miP1a and 773 proteins interacting with miP1b. In total, 303 proteins were in common between miP1a and miP1b. These include among others the CONSTANS-like 4 (COL4) protein, COL9, CONSTANS-like 9 and TOPLESS (Supplementary dataset 2 and 3). This confirms that the miP1a/b microProteins interact with B-Box transcription factors and associate with TOPLESS-like co-repressor proteins *in vivo*. However, we did not identify CO in these pull-down experiments which can be explained by the low abundance of the CO protein. Alternatively, miP1a/b might form different types of repressor complexes involving also other CO-like proteins. In order to find additional interacting proteins with either TOPLESS or JMJ14 which might shed light on the formation of a potential higher order repressor complex, we also generated plants overexpressing FLAG-TPL and FLAG-JMJ14 to co-purify additional interacting proteins (Supplementary dataset 4). Similar to miP1a/b, we also performed parallel immunoprecipitations with Col-0 and transgenic plants expressing FLAG-GFP but this time performed an additional active coal purification step prior injection into the mass spectrometer. Comparative analysis of the four datasets revealed 180 JMJ14-interacting proteins and 145 TPL-interacting proteins that were more than thirty-fold enriched over the background (Fig.3A,B). In total, we identified 82 proteins co-precipitating with JMJ14 and TPL. The JMJ14 dataset includes two NAC transcription factors NAC50 and NAC52 that have previously been identified to interact with JMJ14 (Ning et al., 2015). TPL co-precipitates all other TOPLESS-related (TPR) proteins, supporting recent finding that TPL/TPR proteins form tetramers (Martin-Arevalillo et al., 2017). These examples confirm that our MS-IP strategy identifies *bona fide* JMJ14- and TPL-interacting proteins. We were however surprised to find that none of the previously identified TPL/TPR-interacting repression-domain containing transcription factors (Causier et al., 2012) was present in our dataset. This could indicate that these interactions are either transient or stabilized by additional interacting proteins. Interestingly, the overlap between TPL and JMJ14 interactors includes TPL implying that JMJ14 is part of some TPL repressor complexes. In summary, we provide evidence that a microProtein complex containing both TPL and JMJ14 can form to repress flowering.

**Fig. 3.**
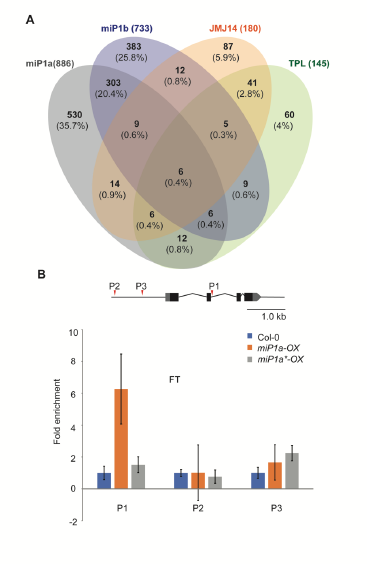
JMJ14 is part of the miP1-TOPLESS repressor complex. **(A)** Venn diagram depicting the number of proteins co-purified with FLAG-miP1a, FLAG-miP1b, FLAG-JMJ14 and FLAG-TPL. Non-specific interactors identified in experiments with either wilt type plants or plants expressing FLAG-GFP have been subtracted. **(B)** Chromatin-immunoprecipitation experiment with wild type and transgenic plants expressing either FLAG-miP1a or FLAG-miP1a*, the latter carrying mutations in the B-Box domain that prevent an interaction with CONSTANS. Gene model depicts the positions amplified by qPCR. Diagram shows a specific enrichment with transgenic plants expressing FLAG-miP1a at position 1.

### The miP1a microProtein interacts with the promoter of *FT*

To confirm that miP1a is part of DNA-binding complex, we performed chromatin-immunoprecipitation experiments with transgenic plants expressing either FLAG-miP1a or FLAG-miP1a*. The latter is a miP1a variant in which all cysteines and histidines of the B-Box zinc finger have been changed to alanine. The miP1a* protein can no longer interact with CO but retains its ability to interact with TPL (Graeff et al., 2016). CO acts as transcriptional activator of *FT* and has been shown to directly and physically interact with the *FT* promoter (Hayama et al., 2017; Song et al., 2012; Tiwari et al., 2010). We used primers amplifying around the previously identified CO-response element (CORE; P3) (Tiwari et al., 2010) plus additional primer pairs amplifying up-stream (P2) and downstream (P1) of the CORE. We did not detect enrichment that would support binding of miP1a around the CORE region but instead detected enrichment indicative of miP1a binding in the second exon of the *FT* gene. No enrichment was observed with transgenic plants expressing FLAG-miP1a* demonstrating that a functional B-Box is required, most likely to associate with a DNA-binding protein that mediates the chromatin-interaction. Interestingly, FLAG-miP1a binding occurs near a recently identified CO-binding site (Hayama et al., 2017), indicating that it is CO which is part of the miP1a DNA-binding complex.

### Mis-expression of *CONSTANS* in the shoot meristem accelerates flowering in *jmj14* mutant plants

*CO* and *FT* are both expressed and active in the leaf vasculature (An et al., 2004) however *CO* is also expressed in the shoot apical meristem (SAM) where *FT* is absent. This could indicate a flowering-independent role of CO in the SAM or a role as a repressor or *FT* expression in the SAM. When expressed from the SAM-specific *KNAT1* promoter, CO is unable to rescue the late flowering phenotype of *co* mutant plants. This contrast findings with *FT*, where expression from the *KNAT1* promoter resulted in very early flowering in the *co* mutant background (An et al., 2004). We noted that besides *CO*, also *miP1a*, *miP1b* (Graeff et al., 2016) and *JMJ14* (Yang et al., 2010) all show a strong expression in the SAM. In agreement with this, we find a very early flowering response when we introduce the *KNAT1::CO* transgene into *jmj14* mutant background (Fig. 4A,B). Also in combination with a mutation in *co*, *KNAT1::CO jmj14 co-2* mutants flower very early supporting the idea that JMJ14 is part of a repressor complex that acts in the SAM to repress *FT* expression.

**Fig. 4.**
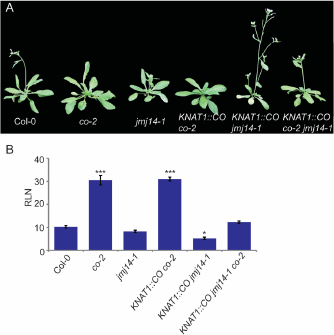
Expression of the *CO* in the meristem of *jmj14* mutants rescues the late flowering phenotype of *co* mutants. **(A)** Picture of representative plants. **(B)** Determination of flowering time by counting the number of rosette leaves at the bolting stage of wild type, *co-2*, *jmj14-1*, *KNAT1::CO co-2* and *KNAT1::CO jmj14-1* and *KNAT1::CO co-2 jmj14-1* mutant plants. *P<0.05, ***P<0.001.

## CONCLUSION

Depending on the age of the plant, the environmental conditions or the tissue, specific transcription factors have been identified that can regulate the transition to flowering. Chromatin-modifying complexes containing polycomb group proteins and diverse histone-modifying enzymes fine-tune the chromatin state of the floral integrator gene *FT* in a plug-and-play fashion (Forderer et al., 2016; Gu et al., 2013; Wang et al., 2014). Here, we provide evidence that microProteins can engage a flowering activator into a floral repressor complex. This repressor complex likely contains additional components that act to modify the chromatin state of *FT*. Mutants affecting the *JMJ14* gene lose their repressive potential, which alleviates flowering. The finding that mutations in *CO* remain late flowering also when *JMJ14* is mutated support a role for CO in this repressive complex. Elucidating these control-circuits in a spatiotemporal fashion will be the next steps in understanding how the balance of activating and repressing complexes triggers developmental transitions.

## EXPERIMENTAL PROCEDURES

### Plant material and growth conditions

Transgenic plants overexpressing miP1a, miP1b and miP1a* are described in (Graeff et al., 2016). The *jmj14-1* mutant corresponds to SALK_135712. For flowering time experiments, seeds were stratified 48h at 4°C, and grown on soil in a plant growth chamber under long day light conditions (16h light /8h dark) at (22°C day /18°C night), or short day light conditions (8h light /16h dark) at (22°C day /18°C night). Flowering time was measured by counting the number of rosette leaves at onset of bolting. Data are expressed as mean +/−SD.

### EMS mutagenesis and growth of Arabidopsis

A seed stock of approximately 1ml homozygote transgenic *35S::FLAG-miP1a* seeds were immersed in 0.025% ethylmethanesulfonate (Sigma) overnight with gentle agitation. These M1 seeds were grown, self-pollinated, pooled and harvested. Approximately 1000 M2 seeds from each original M1 pool were grown in soil under long day conditions to identify early flowering suppressors of miP1a. Suppressors were categorized on the basis of leaf count at flowering. This was defined as plants that flowered with less than or equal leaves at flowering as Col-0, which meant that they flowered significantly earlier when compared to the flowering time of the non-mutagenized parental transgenic plants. They were further characterized by quantification of the *miP1a* mRNA levels by qRT-PCR and protein levels by western blot.

### Identification of mutants and construction of a mapping population

The early flowering *sum1* suppressor plant was backcrossed to the non-mutagenized Col-0 and the late flowering F1 offspring was allowed to self-pollinate. A population of F2 individuals was grown to identify segregating mutants. From 20 early flowering plants one leaf disk of each plant was extracted by a leaf punch and pooled. For the control genome sequencing, five leaf discs each of four miP1a-OX plants were pooled separately. Genomic DNA of these two samples was extracted (DNeasy plant mini kit, QIAGEN). Novogene (Hongkong) prepared libraries and performed sequencing on an Illumina HiSeq4000 (350bp insert size, 100bp paired-end, 7 Gb data).

### Mapping-by-sequencing

More than 95% sequenced reads were mapped by Bowtie2 (v2.1.0) (Langmead and Salzberg, 2012) using the TAIR9 genome assembly and TAIR10 annotation from Phytozome v10.3 (phytozome.org). SNP calling was performed using samtools and BCFtools (v0.1.19) (Li et al., 2009). 1121 (Chr1: 288, Chr2: 233, Chr3: 235, Chr4: 164, Chr5: 201) background corrected EMS-induced SNP markers were identified by SHOREmap v3.2 (Schneeberger et al., 2009) using standard settings. Finally, 591 high quality mutations (quality >= 100, reads supporting the predicted base >= 20) indicated a mapping interval of 2,500 Kb on chromosome 4, containing 10 mutations. The trend line is the average of all SNP allele frequencies in a sliding window (size: 2,500 Kb; step: 100 Kb).

### Gene expression analysis

RNA was extracted from a pool of 12 two weeks old plants from all lines under investigation for gene expression analysis using the Spectrum Plant Total RNA Kit (Sigma-Aldrich). qRTPCR for miP1a, CO and FT was performed as described previously (Graeff et al., 2016).

### Protein purification for mass spectrometry (MS)

Plant tissue from 3-4 weeks old WT, GFP-FLAG-OX, miP1A-OX and miP1B-OX Arabidopsis plants was harvested and flash frozen in liquid nitrogen. The tissue was homogenized and resuspended in SII buffer (100 mM Sodium Phosphate pH 8.0, 150 mM NaCl, 5mM EDTA, 5mM EGTA, 0.1% TX- 100, protease inhibitor (cOmplete^TM^, EDTA-free Protease Inhibitor Cocktail), 1mM PMSF and 1x Phosphatase inhibitors), sonicated and clarified by centrifugation. The protein extract was bound to anti-FLAG M2 magnetic beads (Sigma-Aldrich) for 1hr. Protein bound beads were washed with SII buffer sans inhibitors, followed by washes with 25mM ammonium bicarbonate buffer. The beads were flash frozen with liquid nitrogen prior to downstream analysis.

### Mass Spectrometry parameters

Sample preparation: Proteins bound to anti-FLAG beads were subjected to on-bead digestion as follows: beads were washes 3 times with 10mM Ammonium bicarbonate (pH 7.5-8.0), trypsin was added to each sample and digestion was performed overnight at 37°C. The supernatant was collected and dried by speed vac. The peptides were dissolved in 5% Formic Acid/0.1% Trifluoroacetic Acid (TFA), and protein concentration was determined by nanodrop measurement (A260/A280) (Thermo Scientific Nanodrop 2000 UV-Vis Spectrophotometer). An amount of 0.5ug (5μl) of 0.1% TFA diluted protein extract was injected per sample for LC-MS/MS analysis. LC-MS/MS analysis was performed on a Thermo Scientific Orbitrap Elite mass spectrometer equipped with a Waters nanoAcquity UPLC system utilizing a binary solvent system (Buffer A: 100 % water, 0.1 %formic acid; Buffer B: 100% acetonitrile, 0.1% formic acid). Trapping was performed at 5μl/min, 97% Buffer A for 3 min using a Waters Symmetry^®^ C18 180 μm x 20mm trap column. Peptides were separated using an ACQUITY UPLC PST (BEH) C18 nanoACQUITY Column 1.7 μm, 75 μm x 250 mm (37oC) and eluted at 300 nl/min with the following gradient: 3% buffer B at initial conditions; 5% B at 3 minutes; 35% B at 140 minutes; 50% B at 155 minutes; 85% B at 160-165 min; return to initial conditions at 166 minutes. MS was acquired in the Orbitrap in profile mode over the 300-1,700 m/z range using 1 microscan, 30,000 resolution, AGC target of 1E6, and a full max ion time of 50 ms. Up to 15 MS/MS were collected per MS scan using collision induced dissociation (CID) on species with an intensity threshold of 5,000 and charge states 2 and above. Data dependent MS/MS were acquired in centroid mode in the ion trap using 1 microscan, AGC target of 2E4, full max IT of 100 ms, 2.0 m/z isolation window, and normalized collision energy of 35. Dynamic exclusion was enabled with a repeat count of 1, repeat duration of 30s, exclusion list size of 500, and exclusion duration of 60s.

Protein Identification Database searching: All MS/MS spectra were searched using the Mascot algorithm (version 2.4.0) for un-interpreted MS/MS spectra after using the Mascot Distiller program to generate Mascot compatible files. The data was searched against the Swiss Protein database with taxonomy restricted to Arabidopsis thaliana, allowing for methionine oxidation as a variable modification. Peptide mass tolerance was set to 10ppm and MS/MS fragment tolerance to 0.5 Da. Normal and decoy database searches were run to determine the false discovery rates, and the confidence level was set to 95% within the MASCOT search engine for protein hits based on randomness.

## SUPPLEMENTAL INFORMATION

The paper is accompanied by a supplementary file containing figures S1-S3 and four supplementary datasets containing sequencing and proteomics data.

## AUTHOR CONTRIBUTIONS

VLR, UD, EBP, BS, SH and MG performed the experiments; DS analyzed Illumina sequencing data; TE, MWL and JG performed and analyzed proteomics experiments; SW conceived the research and wrote the manuscript with input from all co-authors

## ACKNOWLEDGEMENTS

We thank George Coupland for providing seeds and the Yale proteomics center and the QBIC center at the University of Tübingen, here the help of Mirita Franz-Wachtel and Boris Maček is specially acknowledged, for proteomics analysis. The work was funded by the Deutsche Forschungsgemeinschaft (WE4281/7-1) and the European Research Council (grant no. 336295) to SW.

---

**Supplemental Figure S1.**
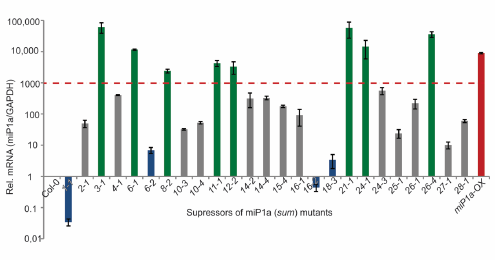
Expression levels of *miP1a* transgene in potential suppressor mutants. Individual plants showing high *miP1a* transcript levels (green bars), comparable to the FLAG-miP1a parental overexpression line (red bar) were isolated for further analysis. Lines with intermediate expression levels (gray bars) and low expression levels (blue bars) were discarded.

**Supplemental Figure S2.**
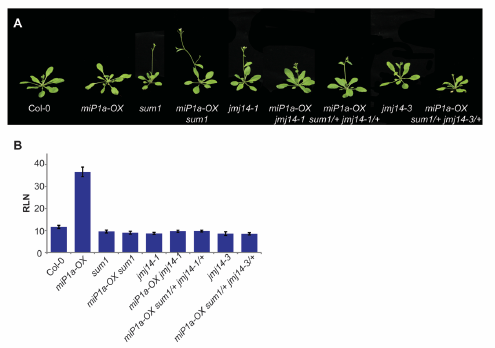
*Sum1* is the phenotype causing mutation. (A) A characterized JMJ14 mutant plant *jmj14-1* was crossed with a late flowering *miP1a-OX* plant. The resultant double homozygote *miP1a-OX jmj14-1* offspring is early flowering. Additionally, the isolated mutant line *miP1a-OX sum1* was crossed into mutants *jmj14-1* and *jmj14-3.* Resultant F1 cross also show an early flowering phenotype. (B) Flowering time counts of the genetic crosses.

**Supplemental Figure S3.**
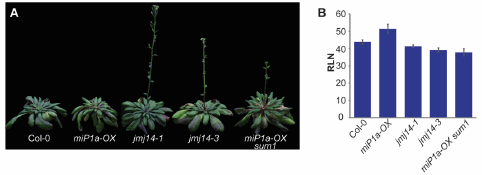
(A) Flowering time of *jmj14* mutants under short day conditions. (8h light / 16h dark) (B) Flowering time counts of the genetic crosses.

## REFERENCES

An, H., Roussot, C., Suárez-Lpez, P., Corbesier, L., Vincent, C., Piñeiro, M., Hepworth, S., Mouradov, A., Justin, S., Turnbull, C., et al. (2004). CONSTANS acts in the phloem to regulate a systemic signal that induces photoperiodic flowering of Arabidopsis. Development (Cambridge, England) 131, 3615–3626

Andres, F., and Coupland, G. (2012). The genetic basis of flowering responses to seasonal cues. Nat Rev Genet 13, 627–639

Causier, B., Ashworth, M., Guo, W., and Davies, B. (2012). The TOPLESS Interactome: A Framework for Gene Repression in Arabidopsis. Plant Physiology 158, 423–438

Corbesier, L., Vincent, C., Jang, S., Fornara, F., Fan, Q., Searle, I., Giakountis, A., Farrona, S., Gissot, L., Turnbull, C., et al. (2007). FT Protein Movement Contributes to Long-Distance Signaling in Floral Induction of Arabidopsis. Science (New York, NY) 316, 1030–1033

Eguen, T., Straub, D., Graeff, M., and Wenkel, S. (2015). MicroProteins: small size-big impact. Trends in plant science 20, 477–482

Forderer, A., Zhou, Y., and Turck, F. (2016). The age of multiplexity: recruitment and interactions of Polycomb complexes in plants. Current opinion in plant biology 29, 169–178

Graeff, M., Straub, D., Eguen, T., Dolde, U., Rodrigues, V., Brandt, R., and Wenkel, S. (2016). MicroProtein-Mediated Recruitment of CONSTANS into a TOPLESS Trimeric Complex Represses Flowering in Arabidopsis. PLoS genetics 12, e1005959.

Gu, X., Wang, Y., and He, Y. (2013). Photoperiodic regulation of flowering time through periodic histone deacetylation of the florigen gene FT. PLoS biology 11, e1001649.

Hayama, R., Sarid-Krebs, L., Richter, R., Fernandez, V., Jang, S., and Coupland, G. (2017). PSEUDO RESPONSE REGULATORs stabilize CONSTANS protein to promote flowering in response to day length. The EMBO journal 36, 904–918

Imaizumi, T., and Kay, S.A. (2006). Photoperiodic control of flowering: not only by coincidence. Trends in plant science 11, 550–558

Jaeger, K.E., and Wigge, P.A. (2007). FT protein acts as a long-range signal in Arabidopsis. Current biology : CB 17, 1050–1054

Langmead, B., and Salzberg, S.L. (2012). Fast gapped-read alignment with Bowtie 2. Nature methods 9, 357–359

Le Masson, I., Jauvion, V., Bouteiller, N., Rivard, M., Elmayan, T., and Vaucheret, H. (2012). Mutations in the Arabidopsis H3K4me2/3 demethylase JMJ14 suppress posttranscriptional gene silencing by decreasing transgene transcription. The Plant cell 24, 3603–3612

Li, H., Handsaker, B., Wysoker, A., Fennell, T., Ruan, J., Homer, N., Marth, G., Abecasis, G., and Durbin, R. (2009). The Sequence Alignment/Map format and SAMtools. Bioinformatics 25, 2078–2079

Martin-Arevalillo, R., Nanao, M.H., Larrieu, A., Vinos-Poyo, T., Mast, D., Galvan-Ampudia, C., Brunoud, G., Vernoux, T., Dumas, R., and Parcy, F. (2017). Structure of the Arabidopsis TOPLESS corepressor provides insight into the evolution of transcriptional repression. Proceedings of the National Academy of Sciences of the United States of America 114, 8107–8112

Mathieu, J., Warthmann, N., Kuttner, F., and Schmid, M. (2007). Export of FT protein from phloem companion cells is sufficient for floral induction in Arabidopsis. Current biology : CB 17, 1055–1060

Ning, Y.Q., Ma, Z.Y., Huang, H.W., Mo, H., Zhao, T.T., Li, L., Cai, T., Chen, S., Ma, L., and He, X.J. (2015). Two novel NAC transcription factors regulate gene expression and flowering time by associating with the histone demethylase JMJ14. Nucleic acids research 43, 1469–1484

Pless, B., Oehm, C., Knauer, S., Stauber, R.H., Dingermann, T., and Marschalek, R. (2011). The heterodimerization domains of MLL-FYRN and FYRC—are potential target structures in t(4;11) leukemia. Leukemia 25, 663–670

Putterill, J., Robson, F., Lee, K., Simon, R., and Coupland, G. (1995). The CONSTANS gene of Arabidopsis promotes flowering and encodes a protein showing similarities to zinc-finger transcription factors. Cell 80, 847–857

Samach, A., Onouchi, H., Gold, S.E., Ditta, G.S., Schwarz-Sommer, Z., Yanofsky, M.F., and Coupland, G. (2000). Distinct Roles of CONSTANS Target Genes in Reproductive Development of Arabidopsis. Science (New York, NY) 288, 1613–1616

Schneeberger, K., Ossowski, S., Lanz, C., Juul, T., Petersen, A.H., Nielsen, K.L., Jorgensen, J.-E., Weigel, D., and Andersen, S.U. (2009). SHOREmap: simultaneous mapping and mutation identification by deep sequencing. Nat Meth 6, 550–551

Searle, I.R., Pontes, O., Melnyk, C.W., Smith, L.M., and Baulcombe, D.C. (2010). JMJ14, a JmjC domain protein, is required for RNA silencing and cell-to-cell movement of an RNA silencing signal in Arabidopsis. Genes & development 24, 986–991

Song, Y.H., Smith, R.W., To, B.J., Millar, A.J., and Imaizumi, T. (2012). FKF1 conveys timing information for CONSTANS stabilization in photoperiodic flowering. Science (New York, NY) 336, 1045–1049

Staudt, A.-C., and Wenkel, S. (2011). Regulation of protein function by microProteins. EMBO reports 12, 35–42

Suarez-Lopez, P., Wheatley, K., Robson, F., Onouchi, H., Valverde, F., and Coupland, G. (2001). CONSTANS mediates between the circadian clock and the control of flowering in Arabidopsis. Nature 410, 1116–1120

Tamaki, S., Matsuo, S., Wong, H.L., Yokoi, S., and Shimamoto, K. (2007). Hd3a Protein Is a Mobile Flowering Signal in Rice. Science (New York, NY) 316, 1033–1036

Tiwari, S.B., Shen, Y., Chang, H.C., Hou, Y., Harris, A., Ma, S.F., McPartland, M., Hymus, G.J., Adam, L., Marion, C., et al. (2010). The flowering time regulator CONSTANS is recruited to the FLOWERING LOCUS T promoter via a unique cis-element. The New phytologist 187, 57–66

Valverde, F., Mouradov, A., Soppe, W., Ravenscroft, D., Samach, A., and Coupland, G. (2004). Photoreceptor regulation of CONSTANS protein in photoperiodic flowering. Science (New York, NY) 303, 1003–1006

Wang, Y., Gu, X., Yuan, W., Schmitz, R.J., and He, Y. (2014). Photoperiodic control of the floral transition through a distinct polycomb repressive complex. Developmental cell 28, 727–736

Yang, W., Jiang, D., Jiang, J., and He, Y. (2010). A plant-specific histone H3 lysine 4 demethylase represses the floral transition in Arabidopsis. The Plant journal : for cell and molecular biology 62, 663–673

